# Diversity and evolution of Radiolaria: Beyond the stars of the ocean

**DOI:** 10.1101/2024.10.02.614131

**Authors:** Miguel M. Sandin, Johan Renaudie, Noritoshi Suzuki, Fabrice Not

## Abstract

Since Ernst Haeckel and the Challenger expedition (1872-1876), Radiolaria have been known as ubiquitous and abundant star-shaped oceanic plankton. Their exquisite biomineralized skeletons have left an extensive fossil record which is extremely valuable for biostratigraphic and paleo-environmental research. In contemporary oceans, there is growing evidence that Radiolaria are significant contributors to marine food webs and global biogeochemical cycles. Here we provide a comprehensive morpho-molecular framework to assess the extant diversity, biogeography and evolutionary history of Radiolaria. Our analyses reveal that half of radiolarian diversity is morphologically undescribed, with a large part forming three hyper-diverse environmental clades, named Rad-A, Rad-B and Rad-C. We suggest that most of this undescribed diversity likely comprises skeleton-less life forms or endosymbionts, explaining their elusive nature. Phylogenetic analyses highlight the need for major revision of high-level Radiolaria taxonomy, including placement of the Collodaria within the order Nassellaria. Fossil calibration of a molecular clock revealed the first appearance of Radiolaria ∼760 million years ago (Ma), the development of the skeleton in the early Paleozoic (∼500 Ma) and the onset of photosymbiotic relationships during the mid to late Mesozoic (∼140 Ma), related to geological periods of oligotrophy and anoxia. The results presented here provide an extensive and robust framework for developing new perspectives on early eukaryotic diversification, paleo-environmental impacts on plankton evolution, and marine microbial ecology in rapidly evolving ecosystems.

## Introduction

Radiolaria are amoeboid planktonic protists that are ubiquitous and abundant in the world’s oceans. Together with Foraminifera they constitute the Retaria lineage, a main branch in the eukaryotic tree of life within the Rhizaria supergroup (1). Radiolaria are historically divided into five groups: Acantharia with a strontium sulphate skeleton, the Polycystinea Collodaria, Nassellaria and Spumellaria with siliceous skeletons and the Taxopodida (encompassing the sole genus *Sticholonche*) with oar-like silicified spines (2). The robust skeleton of Polycystinea preserves very well in sediments, resulting in a continuous fossil record back to the early Cambrian (3, 4) that has been extensively studied by micro-paleontologists, establishing radiolarian taxonomy and evolutionary history from a morphology-based perspective (5–7). This framework provides valuable tools for biostratigraphy (4, 8) and paleo-environmental reconstruction studies (9). In present oceans, Radiolaria are central to ecosystems functioning, representing up to 5% of total biomass in surface waters (10) as well as significantly contributing to the silica cycle (11) and carbon export (10). Radiolaria are considered one of the major carbon transporters to the deep ocean (12).

Global molecular-based diversity surveys of marine plankton have highlighted the ecological roles of specific radiolarian groups and provided access to unexplored morphological diversity. While metabarcoding analyses revealed Acantharia and Collodaria to be highly abundant in the sunlit ocean (13), a broad diversity associated with Spumellaria and Taxopodida appeared to be dominant in deep environments (14). Such environmental diversity from deep environments has been reported since the advent of molecular environmental surveys (15), yet fundamental attributes remain unknown since their morphology, life mode and ecological roles are still a mystery. Current knowledge on Radiolaria diversity suggests that these biogeographical patterns are partially driven by the mixotrophic nature of some Radiolaria species, with Collodaria and Acantharia dominating oligotrophic and eutrophic ocean surface waters, respectively (16). All Collodaria exhibit symbiotic relationships with photosynthetic dinoflagellates (17, 18) and some Acantharia associate with haptophytes (19). By contrast, Spumellaria harbour a variety of photosynthetic symbionts (20), from dinoflagellates to cyanobacteria, prasinophytes, and haptophytes (21–23).

In recent years single-cell molecular characterization combined with morphological identification (*i*.*e*. single-cell barcoding) has contributed to unprecedented exploration of the diversity of Radiolaria from ecological and evolutionary perspectives. Such efforts have resulted, for example, in analysis of co-evolutionary patterns between Acantharia and their symbionts, revealing that this endosymbiosis was most likely triggered by periods of extended oligotrophy in the oceans (19, 24). A well-resolved barcode database additionally allowed an accurate description of collodarian molecular biogeography and diversity at a fine taxonomic level (25). Recent studies have used the extensive knowledge of the fossil record of Polycystinea to time-calibrate phylogenetic trees of Nassellaria (26) and Spumellaria (27), leading to the reinterpretation of ancient fossil groups (28). Further analyses have also demonstrated that orders such as the Entactinaria that were previously considered distinct and homogeneous based on morphological features are actually scattered among Rhizaria (29). Several studies have attempted to unveil high-level relationships between the different groups of Radiolaria (30–32), but the incompleteness of datasets has resulted in inconclusive results.

Here we gather previous morpho-molecular barcoding data along with publicly available environmental sequences (33) and long-read environmental sequencing data (34) into the most comprehensive and non-redundant rDNA-based molecular phylogenetic framework for Radiolaria to date. The extensive fossil record of Radiolaria allowed us to time-calibrate this phylogeny and reconstruct radiolarian diversification contextualized with biotic and abiotic drivers derived from exploration of extant biogeography of Radiolaria through global metabarcoding datasets.

## Results

### Phylogenetic diversity of extant Radiolaria

In total, 658 non-redundant Operational Taxonomic Units (OTUs) of the rDNA gene associated with Radiolaria have been assembled, covering both morphologically described and environmental molecular diversity. Based on comparable phylogenetic distance of sister lineages, bootstrap support and morphological features whenever available, Radiolaria is divided into 6 main groups (**Fig. 1**): Acantharia, Spumellaria, Nassellaria (including the three families of Collodaria), and three highly diverse environmental groups: Rad-A, Rad-B and Rad-C. The latter three groups are composed of environmental sequences with no associated morphological data, with the exception of *Sticholonche*. All six main groups of Radiolaria were highly supported with bootstrap (BS) values above 99, except for Rad-B which exhibited BS values of 91, 87 and 94 for RAxML-CAT, RAxML-ng-GTR+G and IQTree-GTR+G analyses respectively. Spumellaria and Nassellaria together constituted a moderately supported clade (63<BS<76), traditionally known as Polycystinea, sister to Acantharia and the three environmental RAD groups, constituting the Spasmaria group. The latter also formed a moderately supported clade (62<BS<70) with Acantharia and Rad-A as fairly supported sister groups (61<BS<70) and Rad-B and Rad-C as the least supported lineage (60<BS<64).

**Figure 1.**
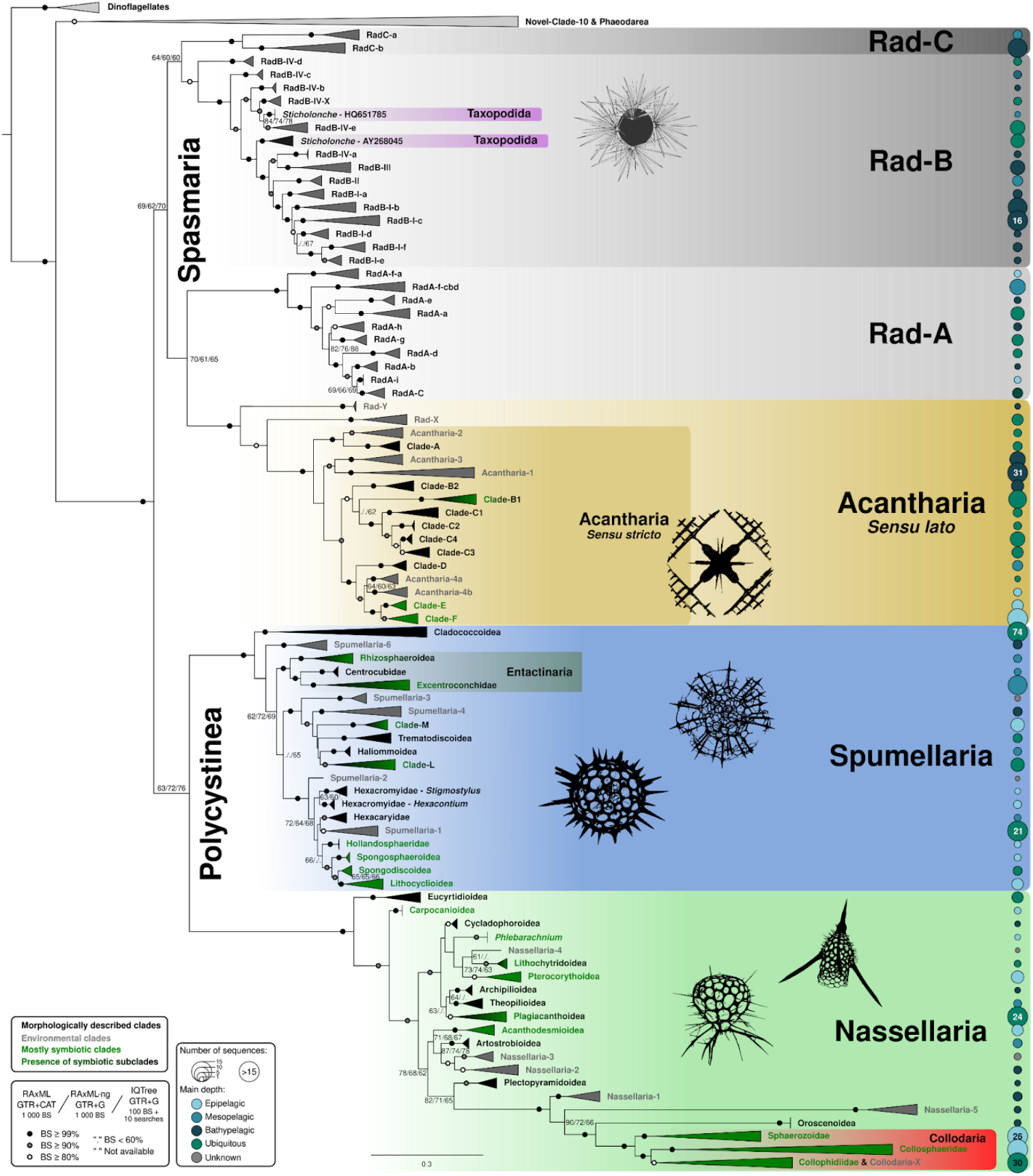
Molecular phylogeny of Radiolaria inferred from the concatenated 18S and 28S rDNA genes (656 taxa and 4920 aligned positions). The tree represents the best scoring tree obtained by RAxML using the GTR+CAT model of sequence evolution. Values at nodes represents RAxML (GTR+Gamma), RAxML (GTR+CAT) and IQTree (GTR+F+R10, best scoring model) bootstrap values (BS; 1000 replicates in RAxML and 100 in IQTree).

Spumellaria and Nassellaria were the most diverse groups with 188 and 157 OTUs respectively, spread among 20 and 21 highly supported clades, based on phylogenetic distance, bootstrap support and morphological characteristics when available. To simplify annotation of different phylogenetic clustering levels, we refer to ‘clade’ as the lowest phylogenetic level and ‘group’ as a monophyletic cluster of clades (*e*.*g*., the clade Plagiacanthoidea within the group Nassellaria, **Fig. 1**). Collodaria formed a strongly supported clade within Nassellaria with 71 sequences, corresponding to nearly half of Nassellaria OTU diversity, spread among only three clades. Out of a total of 20 and 21 clades respectively, both Spumellaria and Nassellaria contained 5 environmental clades each, mostly originating from bathypelagic samples. Acantharia, Rad-B, Rad-A and Rad-C were comprised of a more limited OTU diversity (132, 100, 53 and 19 OTUs respectively), but contained the highest proportion of environmental clades. With 10 out of 15 clades morphologically referenced, Acantharia was the best described group of the Spasmaria lineage. Rad-B comprised 16 clades, with only two sequences morphologically described under the same genus name, despite their relatively large phylogenetic distance (**Fig. 1**; Taxopodida, *Sticholonche*). Rad-A and Rad-C encompassed 10 and 2 clades, respectively, with OTUs composed exclusively of environmental sequences. Within Spasmaria, two orphan environmental clades containing 7 and 2 OTUs (labelled Rad-X and Rad-Y, respectively) were highly supported (BS >99) sister lineages to Acantharia. Given the OTU diversity of Rad-X and Rad-Y, the strong support for them being sister to Acantharia (*sensu stricto*) and their relatively large phylogenetic distance to any other Radiolaria group, here we refer to Acantharia *sensu lato* (comprising Acantharia *s*.*s*. together with Rad-X and Rad-Y) to include all morphologically described and environmental molecular diversity related with confidence to Acantharia.

### Molecular dating of extant Radiolaria

A total of 25 fossil calibration points were used to date the phylogenetic framework shown in **Fig. 1**. According to this analysis, Radiolaria appeared 758 million years ago (Ma; with a 95% Highest Posterior Density -HPD-between 943 and 601 Ma; **Fig. 2**). Spasmaria and Polycystinea then diversified at concomitant times: 652 (HPD: 827-492) Ma and 646 (HPD: 800-538) Ma, respectively. The last event in the Proterozoic was the split between the Acantharia *sensu lato* and Rad-A at 569 (HPD: 742-417) Ma. It is worth noting the variability of the absolute dates of the deep nodes of Radiolaria between BEAST2 (normal distribution prior) and MCMCTree (skew-normal distribution prior) dating, spanning in some cases up to 100 million years of difference (**<u>Fig. S1</u>**). However, the results obtained from BEAST2 and MCMCTree over the Phanerozoic were similar overall. The first appearance of all 6 main groups of Radiolaria described in **Fig. 1** can confidently be stated to have occurred in the Phanerozoic (∼541 Ma or younger). Spumellaria was the first group that diversified 515 (HPD: 521-509) Ma, followed by the separation between Rad-B and Rad-C at 497 (HPD: 694-328) Ma, the appearance of Nassellaria at 464 (HPD: 521-394) Ma and Acantharia *sensu lato* 455 (HPD: 614-318) Ma. Rad-B showed the greatest highest posterior density of all main groups, emerging at 382 (HPD: 547-249) Ma. Acantharia *sensu stricto* diversified at 307 (HPD: 409-220) Ma, at nearly the same time as Rad-A at 311 (HPD: 466-191) Ma. Rad-C was the last main group of Radiolaria to diversify at 265 (HPD: 445-134) Ma. Lastly, Collodaria emerged at 146 (HPD: 194-106) Ma while the other major clades of symbiotic Radiolaria diversified at 128 (HPD: 196-77) Ma within Spumellaria and 78 (HPD: 123-45) Ma within Acantharia. However, establishment of photosymbiosis could have happened at any time between their last common ancestor with non-photosymbiotic relatives and their diversification as a monophyletic clade. In this context the establishment of photosymbiosis occurred no later than 181 (238-132) Ma for Collodaria, 170 (HPD: 240-113) Ma in Spumellaria and 119 (HPD: 180-73) Ma in Acantharia (or 157; HPD: 229-96; Ma if Acantharia-4 turns out to be photosymbiotic).

**Figure 2.**
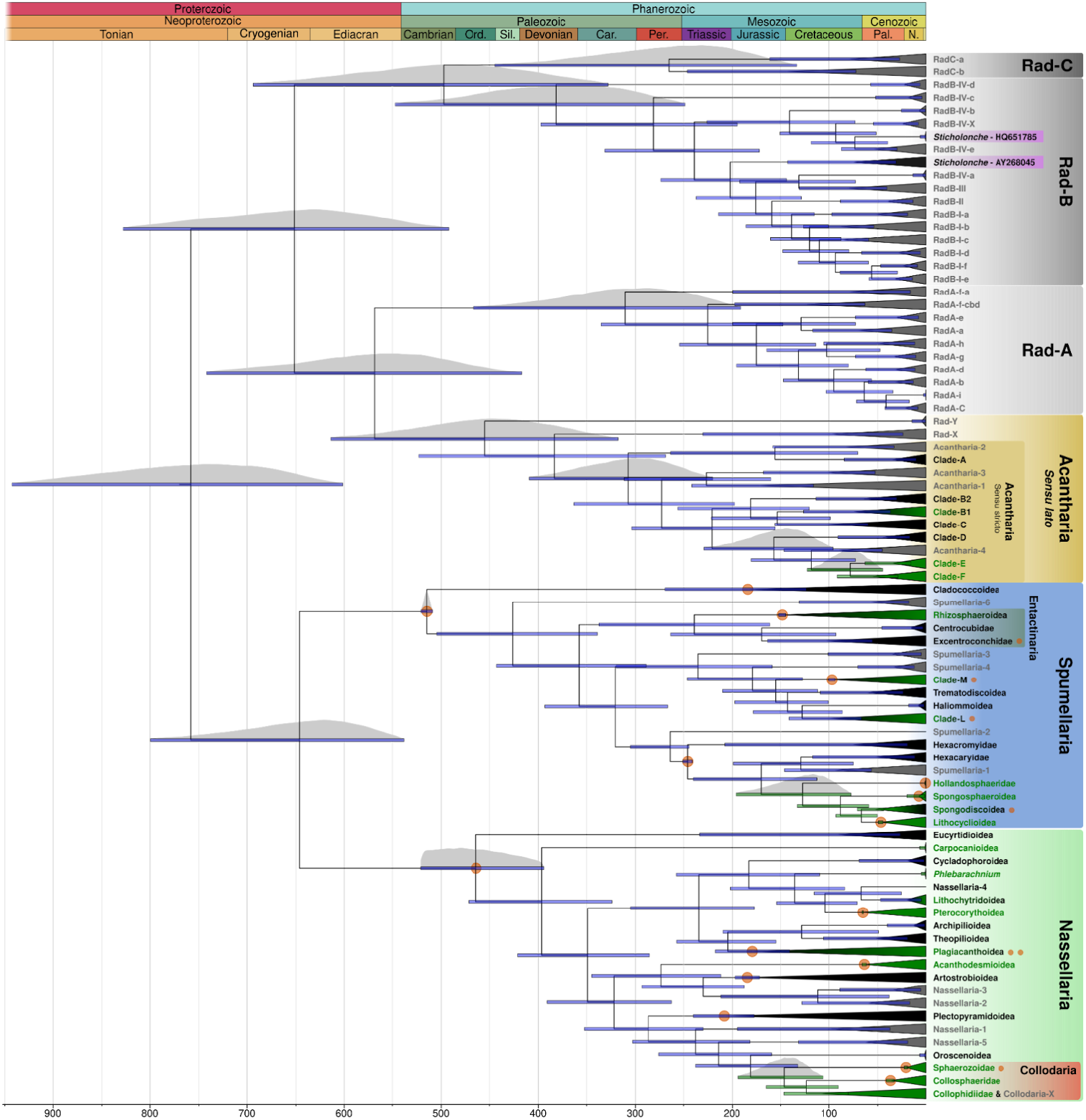
Fossil-calibrated tree (molecular clock) of Radiolaria, based on the alignment matrix used for phylogenetic analyses (**Fig. 1**). Divergence times were inferred in BEAST2 under an optimised relaxed clock model and GTR+G for 4 independent chains of 100M cycles sampled every 1000 steps and a 25% burn in. In total 25 fossil calibrations were used, which are shown with the orange dots behind the blue bars on calibrated nodes or after the clade name if the calibration is within the node (**<u>Fig. S5</u>** shows the prior and posterior distributions of the calibrated nodes). Blue bars indicate the 95% highest posterior density (HPD) intervals of the posterior probability distribution of node ages, shown in most relevant nodes. Black names represent morphologically described clades, grey environmental clades, green symbiotic clades and black to green gradient indicates the presence of symbiotic subclades. Time scale is given in million years ago (Ma).

The concomitant timing for the diversification of Polycystinea and Spasmaria is seen in the stepped Lineages Through Time (LTT) profiles (**<u>Fig. S2</u>**) during the mid to late Neoproterozoic, followed by the appearance of Spumellaria in the Cambrian, Acantharia *sensu lato* and Nassellaria nearly at the same time in the Ordovician and Rad-B in the Devonian. Thereafter, the diversification of lineages was steady until the Permo-Triassic boundary, when a major peak of diversification occurred and the first extant lineages appeared (*i*.*e*.Rhizosphaeroidea, Plectopyramidoidea). After a period of significantly lower diversification, a second peak of origination of extant groups characterized the LTT slopes in the mid Jurassic. Finally, the remaining extant groups diversified in the early Cretaceous, since when diversification remained relatively steady until present.

### Fossil dating of the origin of clades

In a further attempt to reconcile molecular and fossil data and therefore link extant diversity with the fossil record, we applied the recently developed Bayesian Brownian Bridge (BBB) to infer fossil clade origin from fossil data alone. Compared to molecular dating, a number of clades inferred with BBB were older (**<u>Fig. S3</u>** and **<u>Table S3</u>**). This discrepancy might indicate that molecular clades cover a narrower diversity (crown group) than that represented by fossils (total group) which inherently include both extant (crown group) and extinct (stem group) diversity. Clades with a low OTU representation (*i*.*e*. Carpocanioidea, Oroscenoidea, Haliommoidea or Centrocubidae, with only one morphologically-described genus in each clade), were dated much older by BBB than the dates obtained by the molecular clock. On the other hand, clades with high OTU representation and several morphologically-described genera (*i*.*e*.Cladococcoidea, Excentroconchidae or Collosphaeridae; **<u>Table S1</u>**) were dated younger with BBB than with the molecular clock. These clades with high OTU representation are known for their fragile skeleton and therefore their crown diversification might have occurred before that reported in the fossil record. Few clade ages were in accordance between BBB and molecular clock dating, with Plagiacanthoidea and Acanthodesmioidea, both comprised of a wide diversity of both OTU representatives and morphologically described genera, being examples of good agreement between approaches. Lastly, dates of specific last common ancestors between environmental clades sister to morphologically described clades (*i*.*e*. Lithochytridoidea and Nassellaria-4 and Artostrobioidea, Nassellaria-2 and Nassellaria-3) seem to converge among all three different approaches when interpreting such a node as the putative morphologically described clade. Such an agreement when interpreting morphologically described clades along environmental clades might indicate morphologically described clades that have not yet been barcoded molecularly.

### Environmental diversity and biogeographic patterns of the clades identified through metabarcoding

The*Tara* Oceans and*Tara* Polar circle expeditions datasets of the V9 hypervariable region of the 18S rDNA gene were explored to investigate biogeographical patterns of extant Radiolaria. These datasets were chosen due to the extensive metadata associated with each sample, in order to explore main abiotic drivers of extant Radiolaria biogeography. On average, Radiolaria contributed 12.2% (±18.09) to the total eukaryotic community reads, with a minimum of 0.0005% and a maximum of 92.99%. Among Radiolaria, the Collodaria taxa dominated the communities collected by plankton nets in the sunlit ocean, accounting for 94.7, 96.5 and 95.7% of total Radiolaria sequences in the size fractions 5-20 μm, 20-180 μm and 180-2000 μm, respectively (**<u>Fig. S4</u>**). In contrast, the smaller size fractions (*i*.*e*.0.8-3 μm and 0.8-5 μm) that were collected by Niskin bottles at a wider depth spectra, showed a broader diversity of radiolarian sequences. Spumellaria contributed 45.8% and 29.3% to the total Radiolaria abundance in the size fractions 0.8-3 and 0.8-5 μm, respectively. Collodaria nevertheless maintained an important relative abundance after Spumellaria by contributing up to 19.4 and 25.9% in the 0.8-3 and 0.8-5 μm size fractions respectively, followed closely by Rad-B (11 and 24.2% for 0.8-3 and 0.8-5 μm) and Acantharia (22.1 and 10.7% for 0.8-3 and 0.8-5 μm). The different sampling strategies for the different size fractions (net tows vs niskin bottles) accounted for 10.45% of community variability alone.

The biased sampling towards epipelagic waters and specifically surface waters in the datasets used was reflected by a high diversity of colonial Radiolaria metabarcodes in all size fractions with a total of 4234 unique OTUs of the V9 hypervariable region of the 18S rDNA (V9-OTUs; **<u>Fig. S4</u>**). Acantharia was the second most diverse group (557 V9-OTUs), followed by Spumellaria (422 V9-OTUs), Nassellaria (excluding colonial Radiolaria: 375 V9-OTUs), Rad-B (285 V9-OTUs), Rad-A (105 V9-OTUs), Rad-C (86 V9-OTUs) and Rad-X (7 V9-OTUs).

Biogeographical patterns were explored by a Redundancy Analysis (RDA), which allows quantification of community variability explained by environmental parameters. The total 6071 metabarcodes were clustered into 139 unique lineages matching individual OTUs from **Fig. 1**, of which 38 were selected by Escoufier’s method of equivalent vectors as the most significant lineages explaining variability of samples. The RDA had an R-squared of 26.98% (32.48% unadjusted), with 22.11% of the variance explained by the first two axes (**Fig. 3**). The first axis explained 13.94% and separated surface and translucent water samples from mesopelagic samples. Variables such as the optical attenuation coefficient and the optical backscattering coefficient (1/m) were negatively correlated to the first axis, whereas depth and the depth maxima of several different parameters (*i*.*e*.; nitrogen, oxygen, chlorophyll) were positively correlated to the first axis. The second axis explained 8.17% of the variance, and divided equatorial samples with a high sunshine duration (SSD) from samples from high latitudes. From the 38 selected lineages, 12 had little response to environmental parameters in their distribution (contribution < 0.03, data not shown). The remaining 26 lineages showed specific sample preferences according to their high-level phylogenetic grouping, except for Acantharia. Acantharia showed some of the strongest responses to depth variability, with Acantharia-1, Acantharia-3 and Clade-C (*Litholophus* sp.) strongly correlated to deep samples and Clade-E (*Lychnaspis giltschii* and *Coleaspis vaginata*) and Clade-F (*Staurolithium* sp.) correlated to surface samples. Rad-B showed a strong preference for deep waters, in contrast to Rad-A which preferred shallower waters. Colonial radiolarians showed distinct sample preferences compared to other Radiolaria, with strong preferences towards shallow and translucent, or oligotrophic, samples. Lastly, Nassellaria and Spumellaria, at this broad taxonomic resolution, showed no general preferences in terms of depth, but a weak preference towards non-oligotrophic samples.

**Figure 3.**
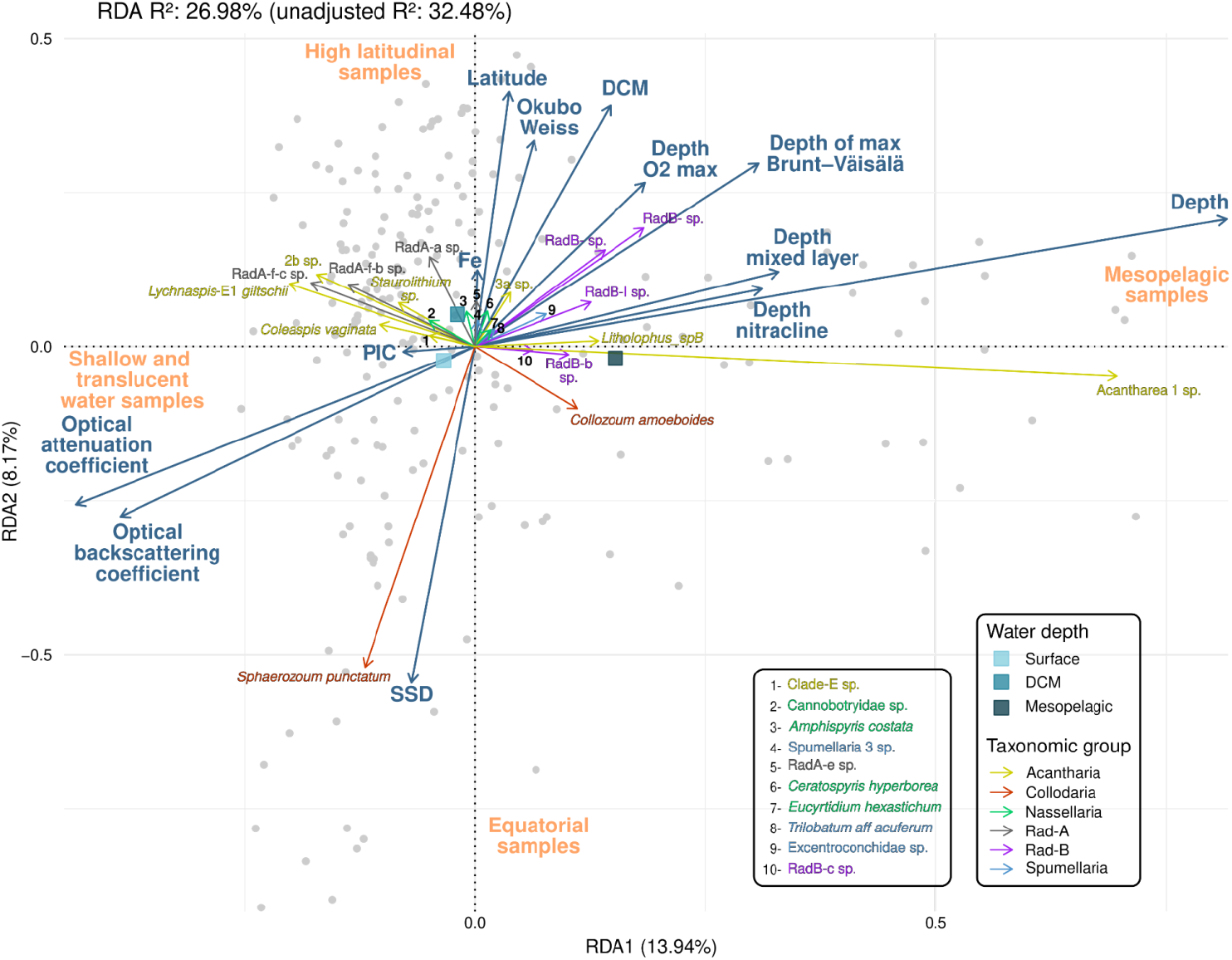
Redundancy analysis (RDA) triplot showing the impact of environmental variables on the distribution of the 38 Escoufier-selected lineages (only those explaining at least 0.03% of the variability are shown) for the smallest size fractions (0.8-3 and 0.8-5μm). The adjusted R-squared of the analysis is of 26.98% (32.48% unadjusted). Abbreviations: SSD: Sunshine duration; PIC: Particulate inorganic carbon; Fe: Iron; DCM: Depth chlorophyll maximum. Text in orange indicates the main environmental parameters of the samples. Note that colonial Radiolaria (Collodaria) lineages have been separated from Nassellaria due to their contrasting environmental preferences.

While Nassellaria and Spumellaria did not show sample preferences in the RDA, our β-diversity analyses identified some lineages associated to the mesopelagic within Spumellaria, such as Cladococcoidea, Rhizosphaeroidea, Centrocubidae and Excentroconchidae (the latter three families constitute the historical group of Entactinaria; **Fig. 1**) as well as within Nassellaria, such as Nassellaria-1 and Cycladophoroidea (**<u>Table S4</u>**). Rad-B strongly dominates the mesopelagic as seen in the RDA. Within Acantharia, Clade-C, Clade-B and Acantharia-1 also showed specificities to mesopelagic water masses, as well as only one lineage within Collodaria corresponding to Collophidiidae. It is worth noting that Collodaria exhibited discrete surface and epipelagic clusters with strong indicator values (**<u>Fig. S7</u>**), in agreement with the RDA results. In terms of latitudinal specificities, several Acantharia lineages associated to the inter-tropical cluster, which corresponded to the clades bearing photosymbionts, as well as Acanthodesmioidea within Nassellaria, Spongosphaeroidea within Spumellaria and several lineages within Collodaria. Spumellaria-1, Spumellaria-6, Rad-A, and *Sticholonche* within Rad-B also showed a strong association to inter-tropical waters. From all lineages analyzed, only two showed specificity towards arctic waters, corresponding to an undescribed lineage within Collodaria (Collophididae) and *Litholopus* within Acantharia (Clade-C).

## Discussion

### On the extant diversity of Radiolaria

Despite extensive morphological characterization of Radiolaria over the last 150 years (35), with more than 7000 species (both extant and extinct) described from the Cenozoic alone (2, 36), here we show that these efforts have merely scratched the surface of understanding actual Radiolaria diversity. Our in-depth analyses call for a major revision of Radiolaria classification at high taxonomic ranks, with division into six major groups corresponding to orders in the Linnaean classification. Acantharia (24) along with three hyper-diverse monophyletic groups that lack morphological description (Rad-A, Rad-B and Rad-C) form the Spasmaria (37), while Nassellaria (as stated herein) and Spumellaria (27) together form the Polycystinea. Collodaria has long been regarded as either a separate order in Polycystinea (1, 2, 28, 38), or within Spumellaria (39). However, our results are in accordance with previous molecular phylogenetic investigations that have strongly supported Collodaria nesting within Nassellaria (29, 30, 32). Here we include the family Oroscenidae (40) and the three families of Collodaria (41) within the Nassellaria and refer to Collodaria as a clade at the superfamily (or suborder) rather than order rank.

Our phylogenetic analysis further revealed that half of Radiolaria clades were composed of environmental sequences only. According to their relative phylogenetic distances and overall OTU diversity, these clades correspond to superfamily or family levels and thus the extent of unknown diversity currently leaves 43 out of 86 (super)families to be described. Some environmental clades (clades exclusively composed of sequences with no morphological identification), such as Spumellaria-4 or Acantharia-2 (**Fig. 1**), are sister to already described groups and have been hypothesized to be already described families that have not yet been genetically barcoded, perhaps due to low sampling effort in deep or extreme environments (24, 27). Similarly, our complementary molecular and fossil analyses suggest that Nassellaria-2 and Nassellaria-3 might be morphologically-described but not yet barcoded diversity within Artostrobioidea and Nassellaria-4 within Lithochytridoidea, which are abundant groups in the deep ocean. In contrast, Rad-B contains only 2 sequences associated with morphologically described specimens, both belonging to the same genus (*Sticholonche*; Taxopodida) on the basis of their oar-like spicules. Considering the phylogenetic distance between these two morphologically described specimens, *Sticholonche* seems likely to represent a species complex (42). The large number of environmental clades related to *Sticholonche* coupled with its unique morphological characteristic might also indicate that spicules are only developed in specific life-stages or environments, and thus that it is only morphologically recognizable on specific occasions.

The rest of the diversity, such as Rad-A and Rad-C, comes from various environments sequenced in different studies and shows clear biogeographical preferences (**Fig. 3**; (43). Specific clades of Radiolaria, such as Spumellaria-1, have been suggested to be endosymbionts (likely parasites) of other Spumellaria based on highly frequent chimeric sequences (27). Similar life-modes have been observed in bathyal benthic Foraminifera which live in the dead tests of other Foraminifera (Xenophyophorea) (44, 45). Given that groups like Rad-A that are highly diversified and abundant in the photic ocean seem to have been overlooked in microscope-based studies, it is possible that they might be endosymbionts of other known, but poorly studied, groups.

Another non-exclusive hypothesis is that part of Radiolaria diversity is actually skeleton-less and therefore has been overlooked, thereby hiding in plain sight. Some naked, amoeboid heliozoans for which no 18S rDNA or other molecular reference data is available, such as Gymnosphaerida (46), have been suggested to be Radiolaria based on ultrastructural features of the axopodial complex. Decelle et al., (24) revealed a progressive evolutionary complexification of the Acantharia skeleton from loose spicules in Clade-A to shell-like skeletons in Clade-E. This led to the suggestion that early diverging environmental clades might lack the characteristic star-shaped skeleton of Radiolaria (47). In this study we found significant novel diversity related to Acantharia that extends the diversity of Acantharia (*sensu lato*) to be characterized. This early branching environmental diversity might not be classical star-shaped Radiolaria, but rather exhibit previously neglected morphologies. The use of long-read sequencing of environmental diversity (34) and single-cell barcoding (48) opens exciting new avenues for holistic analysis of Radiolaria ecology and diversity. In this study we applied conservative thresholds when including molecular diversity in our analyses, meaning it is possible that the molecular diversity of Radiolaria extends far beyond that described here.

### An overview of major evolutionary patterns for Radiolaria

Acknowledging that Radiolaria likely extend beyond the classical star-shaped morphology helps to better understand their early appearance in eukaryotic evolutionary history. According to our molecular clock estimations, extant Radiolaria emerged in the Tonian (∼758 Ma), concomitant with Foraminifera (∼770 Ma; (44, 49)), probably favored by slow oxygenation of the deep ocean before the Sturtian glaciation (50, 51). Biomineralization was already occurring in eukaryotes at this time (52) and has been suggested to have emerged as a detoxification mechanism for specific elements (53, 54). Different lineages of Retaria might have thrived in different environments, and therefore presence of different minerals, as a result of avoiding niche competition between kin lineages, where we see nowadays calcium in Foraminifera, silica in Polycystinea and strontium in Acantharia. The appearance of other small planktonic eukaryotes during the Cryogenian, such as algae and early animals (55), increased prey availability probably contributing to the success of Retaria in the community (56, 57), as seen in the diversification of Spasmaria and Polycystinea between the Sturtian and Marinoan glaciations at ∼650 Ma. By the end of the Neoproterozoic, sterane biomarker indicators for Rhizaria appeared for the first time in relatively high concentrations (58), which could be evidence of the success of the Retaria lineage and its consolidation in Proterozoic communities.

Rising predation pressure from Retaria and other predators has been suggested to have triggered the development of multicellularity in animals (58), as observed in in-vitro experiments (59). In an arms race for survival, several planktonic protist groups developed skeletons and other hard structures to escape predation from the novel macrophage pressure from animals during the Cambrian explosion (60, 61). Such a simultaneous and independent diversification of skeleton shapes explains the independent development of the skeleton in Polycystinea compared to the Spasmaria lineage. Based on the symmetric pattern of the skeleton and our phylogenetic results, the heteropolar skeleton of Nassellaria evolved independently from the concentric skeleton of Spumellaria (7). The most likely ancestor of Spumellaria may have resembled the most primitive Polycystinea forms (Archaeospicularia: (4, 27, 62)) and the extinct genus *Palaeospiculum* could be an early ancestor of the total Nassellaria group, that diversified later into the first heteropolar shapes (Proventocitidae) in the Ordovician. This hypothesis is consistent with the occurrence of the extinct order Albaillelaria with bilateral symmetry and could account for the major proposed contribution of Radiolaria to the food web and silica cycle during the Ordovician Plankton Revolution (63–65). On the other hand, Spasmaria developed the skeleton (found in Acantharia sensu stricto and Sticholonche, Rad-B) later, probably because they were thriving in deep environments, as seen in the preferred environment of the early diverging lineages (**Fig. 3**; (14, 43)). Mesopelagic and bathypelagic waters are known to be diluted environments and the chances of encountering predators are smaller than at the surface. In more recent geological times, only the clades that dwelled towards shallower environments (*i*.*e*.Clade-E and Clade-F) might have developed a more elaborate skeleton compared to the loose spicules from Clade-B and Clade-C.

The Mesozoic witnessed the first diversification of extant clades (**Fig. 1**), such as Plectopyramidoidea (6, 8, 26, 27). This Era was the most oligotrophic period of the Phanerozoic (66), and together with long periods of oceanic anoxia (67) the Mesozoic was presumably a challenging environment for heterotrophic eukaryotes. It was at this time that some of the most abundant extant photosynthetic planktonic groups appeared, including dinoflagellates (68), coccolithophores (69) and diatoms (70, 71). The Mesozoic fossil record shows a drastic change in Polycystinea morphologies with many novel shapes (72), which is reflected in the sudden bursts of diversification in the LTT slopes (**<u>Fig. S2</u>**). During the late Jurassic-early Cretaceous, Spumellaria experienced an intense diversification of flat shapes (27) and Nassellaria developed the characteristic gelatinous matrix of Collodaria and *Phlebarachnium* sp. (73). The gelatinous matrix of Nassellaria has been suggested to be an independent evolutionary innovation to cope with oligotrophy in the oceans (73), further supported by the contrasting biogeographical patterns of Collodaria compared to radiolarian relatives as observed in our analyses.

Of all of the evolutionary innovations to cope with the oligotrophic conditions in the Mesozoic, probably the most widespread in Radiolaria was the establishment of symbioses with photosynthetic algae (*i*.*e*. photosymbiosis). Photosymbiotic radiolarians are most commonly associated with dinoflagellate symbionts (historically known as ‘*zooxanthellae*’), found throughout Polycystinea (17, 18) and in Acantharia Clade-B1 (74). According to our phylogenetic reconstruction, the onset of photosymbiosis occurred independently in different lineages of Radiolaria, mostly from the mid-Jurassic and through the Cretaceous (**Fig. 2**, green bars). Contemporarily with Radiolaria, Foraminifera and corals also established photosymbioses with dinoflagellates (75, 76). The number of monophyletic clades at (super)family rank within both symbiotic dinoflagellates and hosts along with theur independent diversification patterns suggest that the establishment of photosymbiosis witnessed several extinction-speciation events, as reflected in the fossil record of corals as well (77). The vast oceanic environment probably limits the encounter of host and symbiont, threatening the fitness of the host at geological time scales and favoring a low host-symbiont specificity. This could explain the diversity of photosynthetic symbionts found in Spumellaria and Acantharia Clade-B1 (20), from dinoflagellates to cyanobacteria, prasinophytes, and haptophytes (21–23). In contrast, Acantharia clades E and F established an exclusive endosymbiosis with the haptophyte *Phaeocystis* (19), where the Acantharia host is capable of dramatically increasing the metabolic productivity of the symbionts’ photosynthetic machinery (78). Collodaria symbionts also show more voluminous plastids than their free-living state (79) and Collodaria have never been reported without dinoflagellate symbionts. Therefore, despite the theoretical lower fitness of a high specificity between host and symbiont at a macro-evolutionary level, our analyses show the undoubted ecological success of extant Acantharia and Collodaria.

The ecological success of Collodaria is only reflected in the fossil record from the middle Cenozoic (∼36 Ma; (6, 36)), although it is still debated when the first representatives of Collodaria appeared in the fossil record (Cenozoic, Triassic or Paleozoic; (6)). Our molecular analyses confidently showed that Collodaria diversified from ancient Nassellaria forms, yet given the divergent morphology of Collodaria, their photosymbiosis dependency and the patchy fossil record, Collodaria most likely underwent a dramatic morphological innovation due to several speciation-extinction events. Other Nassellaria clades sister to Collodaria, such as Oroscenoidea and Acanthodesmioidea (Lineage II; (26)), also show a contrasting morphology compared to the classic cone-shaped of Nassellaria, probably mirroring similar evolutionary events, yet difficult to relate to fossil groups. Recent efforts to link extant Radiolaria diversity with fossils across the Cenozoic (27) allowed the calibration of the molecular clock in this study and thus the unprecedented reconstruction of the evolutionary history of Radiolaria. However, we will only achieve an accurate contextualization with fossil groups and fill the gaps in Radiolaria evolutionary history when all morphologies will be linked back to the first fossil representatives in the Cambrian. Here we connected Radiolaria diversification with global geological and environmental changes to gain insights into their evolutionary history. It is uncertain how Radiolaria will adapt to rapidly changing conditions, since plankton may not be able to shift their geographic distribution at a pace allowing to mitigate climate change (80). Based on the evolutionary history suggested herein and the current expectations of plankton adaptation to global warming (81, 82), we expect Collodaria and Acantharia to increase their diversity towards oligotrophic and high latitude waters, respectively. Although Radiolaria’s fate is uncertain due to the current rate of climate change, the comprehensive framework presented here provides the means to track future community shifts at an unprecedented scale.

## Material and Methods

### Taxonomic curation of environmental sequences associated to Radiolaria

Recent studies on Acantharia (24), Collodaria (41), Nassellaria (26) and Spumellaria (27) have resulted in the detailed morphological description and phylogenetic annotation of nearly 400 18S rDNA (and 28S rDNA) sequences. In addition, a wide diversity of environmental 18S rDNA sequences from environmental samples (mostly clone libraries) have been reported to be closely related to Radiolaria. Using the recently published morpho-molecular framework of Radiolaria, we phylogenetically curated all publicly available 18S rDNA sequences associated to Radiolaria using a step-wise phylogenetic approach (for details see ‘MiguelMSandin/radiolaria/curation_pipeline.md‘). Briefly, a backbone phylogenetic tree of Radiolaria was built including only high quality sequences from morphologically identified specimens, then environmental sequences reported in (24, 26, 27, 41) were added, followed by additional sequences retrieved from public databases. In total, 4556 18S rDNA sequences were phylogenetically curated and uploaded to the Protist Reference Ribosomal (PR2) database from v4.14.0 (33).

### Dataset assembly

A total of 149 concatenated 18S and partial 28S (D1+D2 region) rDNA sequences associated to morphologically identified Acantharia (24), Collodaria (41), Nassellaria (26, 40, 73) and Spumellaria (27) were manually selected. Two additional 28S rDNA sequences belonging to Nassellaria were generated in this study (following (26) and deposited in NCBI with the accession numbers PP648172 (corresponding to *Callimitra emmae*) and PP648173 (corresponding to *Litharachnium tentorium*). These 151 sequences were selected as reference sequences representative of each morphologically described clade based on their maximum length and quality (**<u>Table S1</u>**). 18S rDNA sequences from the Protist Reference Ribosomal (PR2) database v4.14.0 (33); curated herein) were then added to the dataset in order to include uncharacterized environmental diversity. In order to reduce redundancy, the PR2 database was clustered into 97% similarity Operational Taxonomic Units (OTUs) using mothur v1.44.11 (83) after aligning the sequences with MAFFT v7.313 (84). The centroid sequence was taken as representative of each of the 293 OTUs in the PR2 database and OTUs 100% identical to the previously selected sequences were removed, resulting in 269 97% OTUs selected from the PR2 database. In addition, a recently generated long-read metabarcoding dataset spanning the near full rDNA operon (18S-V4 to V9-and 28S rDNA; (34); doi: 10.6084/m9.figshare.15164772) was included to extend rDNA molecular diversity and phylogenetic information. This dataset was sequenced using the Pacific Bioscience Circular Consensus Sequencing workflow and will therefore be referred to as the PacBio dataset from here forward. From the 1044 OTUs assigned to Radiolaria, only those with at least 100 reads and non-identical to an already selected sequence (reference + PR2) were considered, resulting in an additional 254 OTUs of the near-full length rDNA. The conservative threshold for including only OTUs with at least 100 reads was chosen to ensure the inclusion of biologically meaningful sequences and therefore minimize the chances of dealing with PCR/sequencing errors, chimeric sequences or other artefacts. In total, 674 non redundant 18S and 28S rDNA sequences were assembled, representing the most comprehensive molecular diversity of Radiolaria to date.

### Phylogenetic analyses

The 674 assembled sequences were combined with 15 sequences manually selected from Phaeodaria (85), 5 sequences of Novel-Clade-10 and 8 sequences of dinoflagellates as outgroups (**<u>Table S1</u>**). The resulting 702 sequences were split in two datasets corresponding to the 18S rDNA and the 28S rDNA genes. Both datasets were independently aligned with MAFFT v7.511 (84) using two different alignment algorithms (FFT-NS-i and L-INS-i) and 1000 refinement cycles. Aligned datasets were independently trimmed with trimAl v1.4.1 (86) using a 5% threshold and subsequently both genes were concatenated with the in-house script ‘fastaConcat.py‘. Phylogenetic analyses were performed over the two aligned and concatenated datasets resulting from the two different alignment algorithms applying three different maximum likelihood approaches; in RAxML v8.2.12 (87) under the nucleotide substitution model GTR+CAT over 1000 bootstraps each, in RAxML-ng v1.1 (88) under the model GTR+F0+G4m over 1000 bootstraps each and in IQ-Tree v2.0.3 (89) under the model GTR+F+R10 (chosen based on the highest Bayesian Information Criterion from modelFinder) over 100 bootstraps and 10 independent runs. Resulting phylogenetic trees led to the manual removal of up to 16 sequences (6 OTUs from PR2 and 10 OTUs from the PacBio dataset) due to their long branches and dubious positions (**<u>Table S1</u>**). After removal of such sequences, filtered datasets were aligned, trimmed and concatenated as previously described. The 18S rDNA dataset had 675 sequences and 2762 and 2698 positions for the FFT-NS-i and L-INS-i alignment algorithms respectively. The 28S rDNA dataset had 360 sequences and 3859 and 3794 positions for the FFT-NS-i and L-INS-i algorithms respectively. After trimming, the 18S rDNA dataset had 2149 and 2056 positions and the 28S rDNA 2885 and 2846 positions for the FFT-NS-i and L-INS-i algorithms respectively. Final concatenated datasets had 686 sequences and 5034 and 4902 positions for the FFT-NS-i and L-INS-i respectively. Final phylogenetic analyses were run with three different approaches (GTR+F+R10 was the nucleotide substitution model always chosen by the highest Bayesian Information Criterion from modelFinder) as previously described. Both alignments resulted in nearly-identical topologies and therefore the L-INS-i alignment was kept for downstream analyses (available along with the 18S and 28S rDNA alignments before trimming in Zenodo, doi: 10.5281/zenodo.13286957 and at the gitHub repository github.com/MiguelMSandin/radiolaria).

### Molecular dating

The resulting dataset (aligned with the L-INS-i algorithm with MAFFT) from the previous phylogenetic reconstruction step was used for molecular clock analyses implemented in MCMCTree from the PAML v4.10.6 package (90) and in BEAST2 v2.7.4 (91). In total, 23 fossil calibrations were used (**Table 1**) according to (26–28). A detailed justification of the chosen calibrations can be found in **<u>File S1</u>**. The root node was calibrated between 1100 and 2200 million years ago (Ma) after the latest molecular clock results on the diversification between Alveolates and Rhizaria (92, 93) using a uniform distribution and allowing a significant uncertainty due to the lower representation of Alveolata molecular diversity in our dataset. When using MCMCTree, fossil calibrations followed skew-normal distribution where 97.5% probability distribution falls within the minimum and maximum calibrations, estimated in the R package ‘MCMCtreeR’ (94).

**Table 1.**
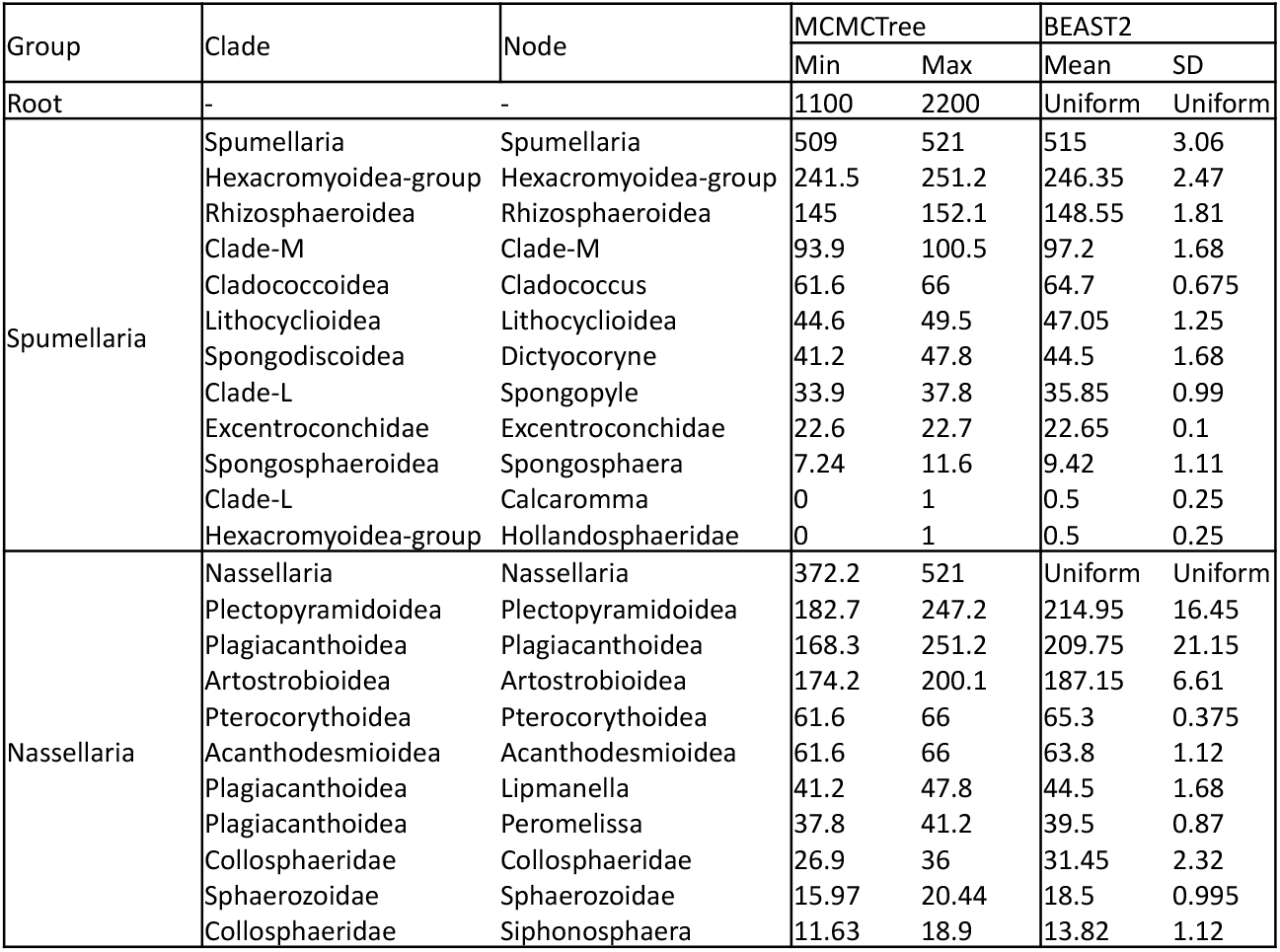
Fossil calibrations used for molecular dating indicating the fossil group, the calibrated node, the minimum (Min) and maximum (Max) appearance time and the mean (Mean) and Standard Deviation (SD). Minimum and Maximum dates were used along a skew-normal distribution. Mean and standard deviation were used along a normal distribution. The root node was always calibrated with a uniform distribution.

Molecular dating with MCMCTree was performed in four steps (adapted from (95) over the resulting phylogenetic tree with the highest logLikelihood (RAxML GTR+CAT), briefly: (i) fossil calibrations and model parameters were checked by sampling the prior on node ages; (ii) maximum likelihood branch length estimators and other model parameters were estimated under the GTR+Gamma (4 categories) nucleotide substitution model; (iii) Markov Chain Monte Carlo (MCMC) sampling of the posterior and rates was run under an independent rate clock model and the birth-death speciation process over 4 independent analyses of 1 million cycles each sampled every 2 states and an initial burn in of 250000 states (25%); and (iv) after checking for convergence and ensuring all parameters and nodes reached at least 200 Effective Sample Size (ESS) in the R package ‘coda’ (96), the 4 independent runs were merged with the in-house script ‘mcmcTreeFilesCombiner.py‘ and the final tree was generated in MCMCTree. When using BEAST2, fossil calibrations followed a relaxed normal distribution such that the 95% probability distribution falls within the maximum and minimum appearance time.

Molecular dating with BEAST2 was performed under the Optimized Relaxed Clock (97) model to avoid assumption of substitution rate correlation between sister clades and a GTR+Gamma (4 categories) model of nucleotide substitution and a Birth-Death model of speciation (98). The resulting tree from sampling the prior in MCMCTree was used as the initial tree to avoid ‘-Infinity’ likelihood issues at the root node (therefore allowing initialization of the MCMC sampling). The tree topology was fixed over the MCMC sampling by setting corresponding parameters to 0, and the MCMC chain was run over 100 million generations sampled every 1000 states on 4 independent runs (remaining operators were left as default). After completion, the 4 independent runs were combined in LogCombiner v2.7.4 with a 25% burn-in and the final tree was summarized in TreeAnnotator v2.7.3. Final control and XML files used to run the analyses stated here are available in the gitHub repository github.com/MiguelMSandin/radiolaria and in Zenodo (doi: 10.5281/zenodo.13286957).

### Molecular clock post-analyses and fossil dating

In order to better integrate the evolutionary history of Radiolaria from the different analyses, we measured a proxy of the diversification over time by a Lineages Through Time (LTT) analysis upon the dated trees obtained from the previous step. In addition, we inferred the age of morphologically described and fossilizable clades by a Bayesian Brownian bridge (BBB; (99)) on the basis of the present diversity and the known fossil record. Fossil data for the BBB analysis was compiled from PBDB (for Mesozoic and Paleozoic data; (100)) and Neptune (for Cenozoic data; (101) databases and living data from the mikrotax database (**<u>Table S2</u>**; (102). Only formally described species (i.e.; excluding open nomenclature taxa) were considered to avoid possible redundancy. Highest posterior density was inferred for the 5% and 95% intervals as well as the median for every analysed clade, over all 25000 iterations of BBB, excluding a burn-in of 5000, using base R.

### Metabarcoding analyses

Global Radiolaria biogeography was analysed from data from the worldwide Tara Oceans (2009-2013) and Tara Polar Circle (2013) expeditions, processed at the OTU level and publicly available at 10.5281/zenodo.3768510 (103). OTUs of the V9 rDNA assigned to Radiolaria were extracted and taxonomically reassigned against the curated version of PR2 (v4.14.0) presented herein. From the 29206 OTUs assigned to Radiolaria, those with an identity score equal to or higher than 90% against a reference sequence and with a total abundance of at least 10 reads and present in at least 2 samples were considered. These filters resulted in the selection of 6071 OTUs for downstream analyses.

Contextual data was assembled by integrating features from the mesoscale (doi: 10.1594/PANGAEA.875582), sampled depth (doi: 10.1594/PANGAEA.853810) and sample collection (doi: 10.1594/PANGAEA.875580). Specific details can be found in (104). The final non-redundant environmental table used in this study can be found in the gitHub repository github.com/MiguelMSandin/radiolaria and in Zenodo (doi: 10.5281/zenodo.13286957).

In order to give a direct biological meaning to the OTUs and diminish biases intrinsic to short-reads (48, 105), filtered OTUs were grouped together at the finest phylogenetic level into lineages (corresponding to species in the PR2 database ranking) resulting in 139 lineages and read abundances were Hellinger transformed. The effect of the arbitrary threshold selected for the different size fractions (0.8-3, 0.8-5, 5-20, 20-180, 180-2000 μm) was tested using a PERMANOVA test along the resulting 139 lineages over 10000 permutations and a Jaccard distance. Final variability explained by size fractions alone was 10.45%. In order to remove this effect from global analysis, the smallest and most diverse (see results) size fraction (pico-nano: 0.8-3 & 0.8-5 μm) was used for statistical analyses to infer Radiolaria biogeography through a redundancy analysis (RDA). From the 139 lineages belonging to the smallest size fraction, 38 were pre-selected by Escoufiers vector as representatives of the community. A two directional stepwise model selection based on AIC (using a maximum model followed by null model) was used to estimate the best formula and conduct the RDA over 100 million steps, 10000 permutations and 3 independent replicates (to account for stochasticity). Analyses were performed in R v4.3.1 with the packages ‘vegan’ v2.6-4 (106) and ‘pastecs’ v1.3.21 (107) and plotted with ‘ggplot2’ v3.4.2 (108).

To further explore the biogeography of Radiolaria, indicator values were calculated (with the function indval from the R package labdsv; (109) for all taxonomic lineages and inferredβ-diversity patterns. Theseβ-diversity analyses are targeted to identifying lineages that are specific to a defined set of samples. To do so, both depth and latitude were discretized in three independent clusters, each corresponding to surface (0-5m deep), epipelagic (5-200m deep; surface and epipelagic were separated due to the disproportional amount of samples at 5m depth), and mesopelagic (200-800m deep); and intertropical (within the tropics), temperate (in between the tropics and the Arctic and Antarctic polar circles), and arctic (above the Arctic polar circle, since there are no samples in the Antarctic), respectively.

## Acknowledgments

This work was supported by the IMPEKAB ANR 15-CE02-0011 grant and the Brittany Region ARED C161520A01. MMS was partially supported by a postdoctoral fellowship from the *Beatriu de Pinós* programme of the Government of Catalonia’s Secretariat for Universities and Research of the Generalitat de Catalunya Economy and Knowledge (grant number: 2021BP00068). We acknowledge the MOOSE program (Mediterranean Ocean Observing System for the Environment) coordinated by CNRS-INSU and the Research Infrastructure ILICO (CNRS-IFREMER). We are also grateful to the CNRS-Sorbonne University ABiMS bioinformatics platform (http://abims.sb-roscoff.fr) for providing computational resources. We would like to acknowledge Dr. Nicolas Henry for advice on metabarcoding analyses, Dr. Cedric Berney for phylogenetic discussions and Dr. Ian Probert for comments on earlier versions of this manuscript.

## Supplementary Material

**Figure S1**: Fossil-calibrated tree (molecular clock) of Radiolaria, based on the alignment matrix used for phylogenetic analyses (**Fig. 1**). Divergence times were inferred in MCMCTree under an uncorrelated rates clock model and GTR+G for 12 independent 20000 length chains sampled every 2000 steps. In total 25 fossil calibrations were used, which are shown with the orange dot behind the blue bars on calibrated nodes or after the clade name if the calibration is within the node (**<u>Fig. S6</u>** shows the prior and posterior distributions of the calibrated nodes). Blue bars indicate the 95% highest posterior density (HPD) intervals of the posterior probability distribution of node ages, shown in most relevant nodes. Black names represent morphologically described clades, gray environmental clades, green symbiotic clades and black degraded to green indicate presence of symbiotic subclades.

**Figure S2**: Lineages Through Time (LTT) analysis based on the molecular clock results shown in **Fig. 2** and **<u>Fig. S1</u>**. The y-axis represents the number of lineages (N) expressed in logarithmic (base e) scale (Ln(N)) and the x-axis represents the time in millions of years ago (Ma). Horizontal gray bars represent the 95% Highest Posterior Density (HPD) of molecular clock estimates.

**Figure S3**: A reconstructed tree based on the Bayesian Brownian Bridge (BBB) dating of fossil clade appearance, compared with those obtained by the molecular clock. Grey dotted lines represent uncertainty in the dates of diversification. Box in the left lower corner represents a schematic interpretation of the contrasting results between BBB and molecular clock analyses, highlighting the different nodes where the conflict might appear depending on the sampled extant diversity.

**Figure S4**: A tree map showing the metabarcode count (upper row) and the abundance (lower row) per main group of Radiolaria. The area represents the proportion of total number of unique metabarcodes or their abundance affiliated to the specific taxonomic entity. Note that Collodaria is treated independently than Nassellaria due to historical reasons and their great contribution.

**Figure S5**: Prior and posterior distributions of the calibration nodes used in BEAST2. Numbers after the node names represent the maximum and minimum fossil appearance times.

**Figure S6**: Prior and posterior distributions of the calibration nodes used in MCMCTree. Numbers after the node names represent the maximum and minimum fossil appearance times.

**Figure S7**: Indicator values of the different discrete clusters stacked for the different metabarcoding lineages ordered by decreasing total indicator value from **table S4**.

**Table S1**: List of ribosomal DNA sequences used in this study.

**Table S2**: A table with fossil diversity per 5 million years time interval of the Polycystinea group used in the Bayesian Brownian Bridge analyses.

**Table S3**: Summary of the Bayesian Brownian Bridge results showing the 0%, 5%, 25%, 50% 75%, 95% and the 100% highest posterior density.

**Table S4**: Indicator values of the 139 metabarcoding lineages for depth (with three discrete cluster: surface: 0-5m, epipelagic: 5-200m and mesopelagic: 200-800m) and latitudinal distribution (intertropical, temperate and arctic). Only significant (p-value < 0.05) indicator values are shown. Strong indicator values are labeled in the column titled ‘strong_indicator’ if the indicator value is above 0.5 and the p-value < 0.01.

**File S1**: A justification of the chosen fossil calibrations.

