## Supplementary figures and images for "Diversity and evolution of Radiolaria: Beyond the stars of the ocean"

### Figure S1

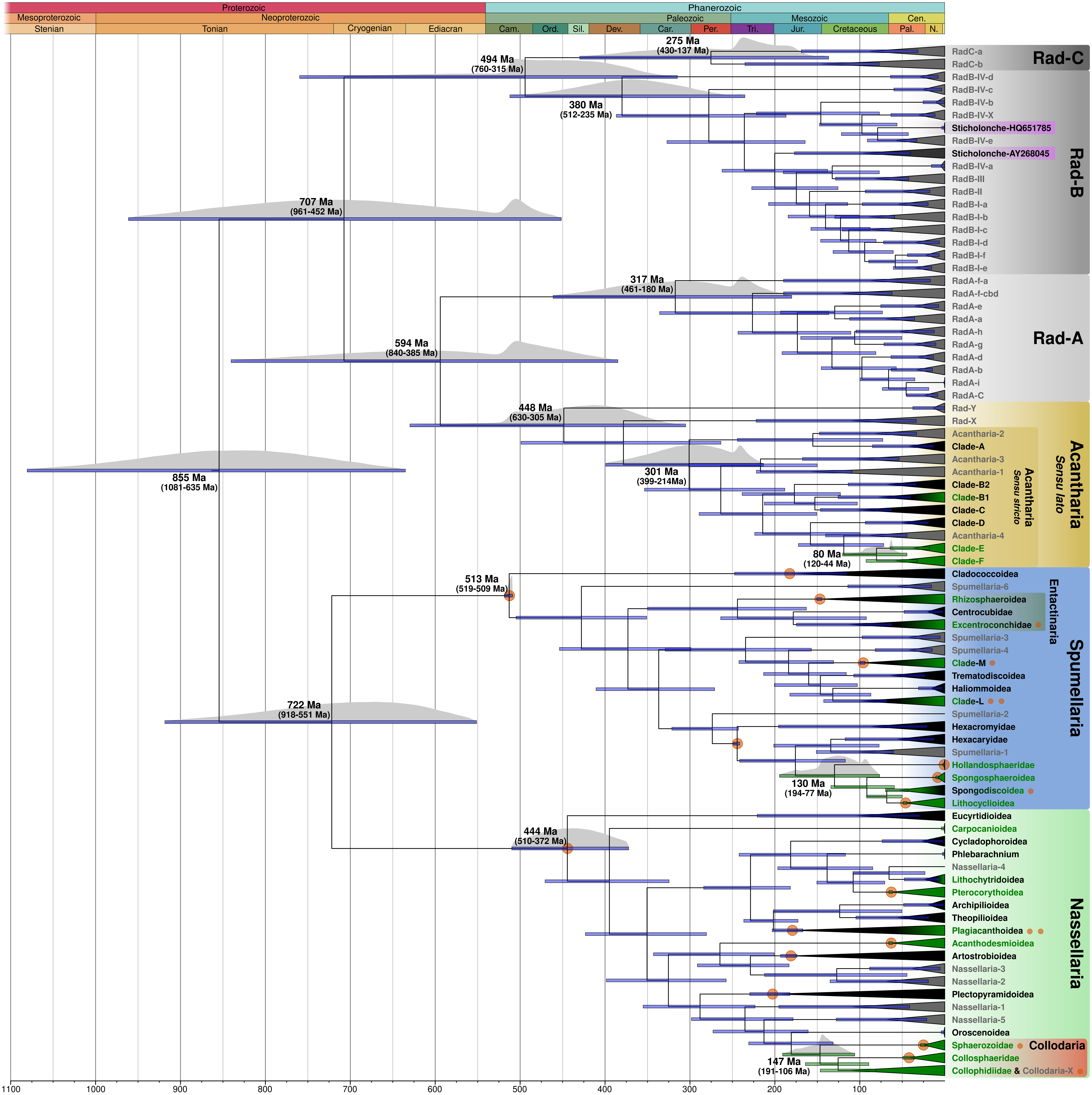

### Figure S2

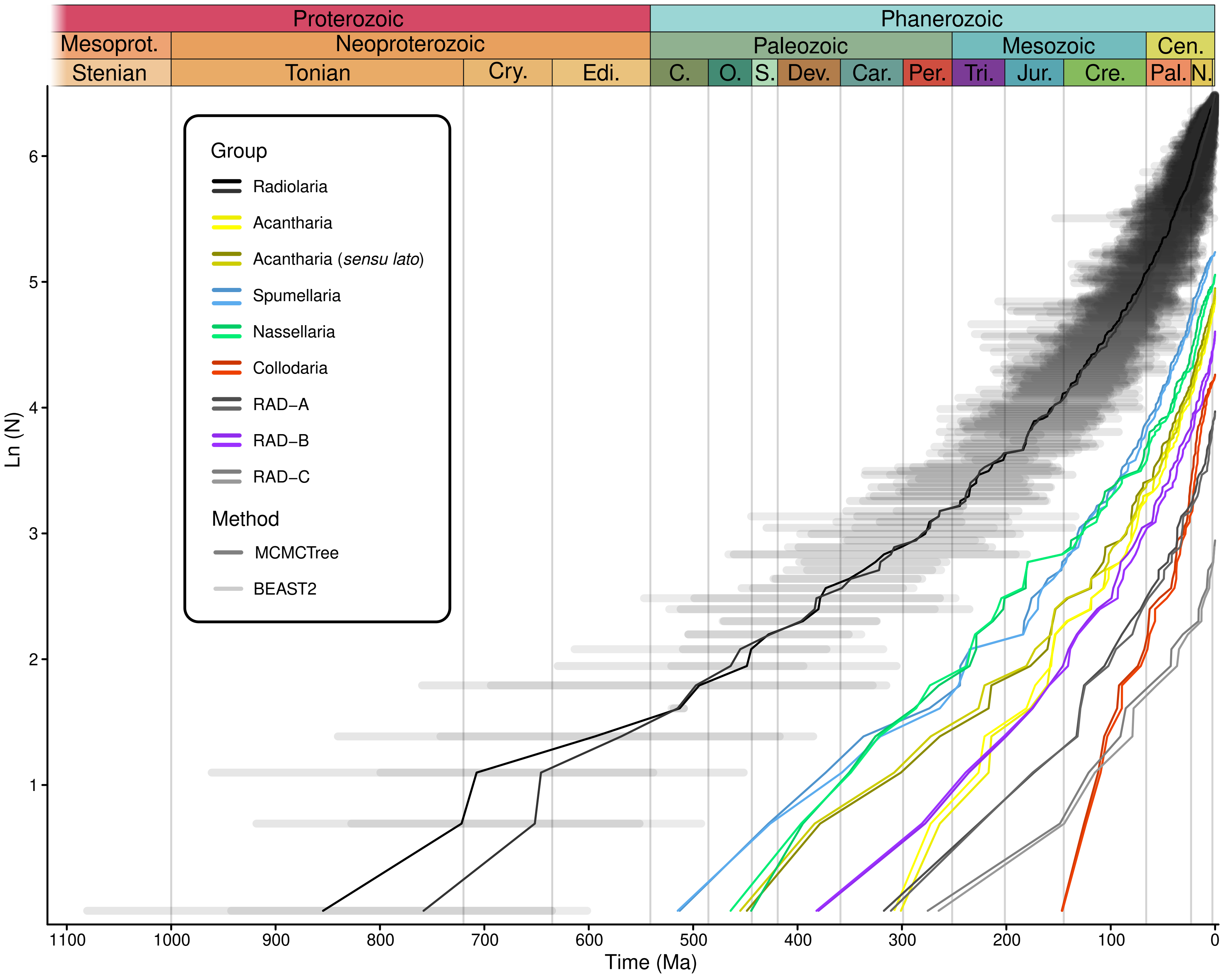

### Figure S3

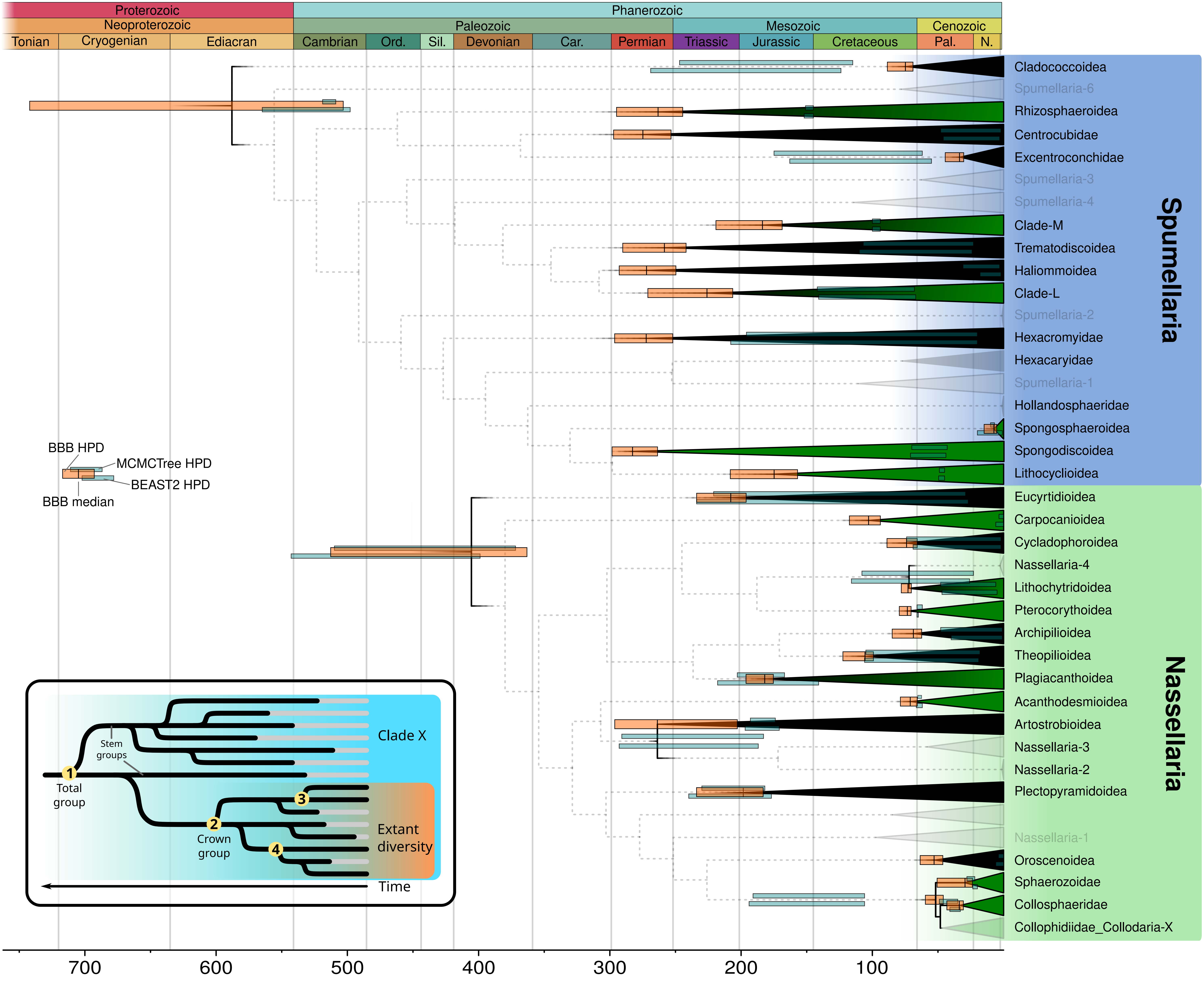

### Figure S4

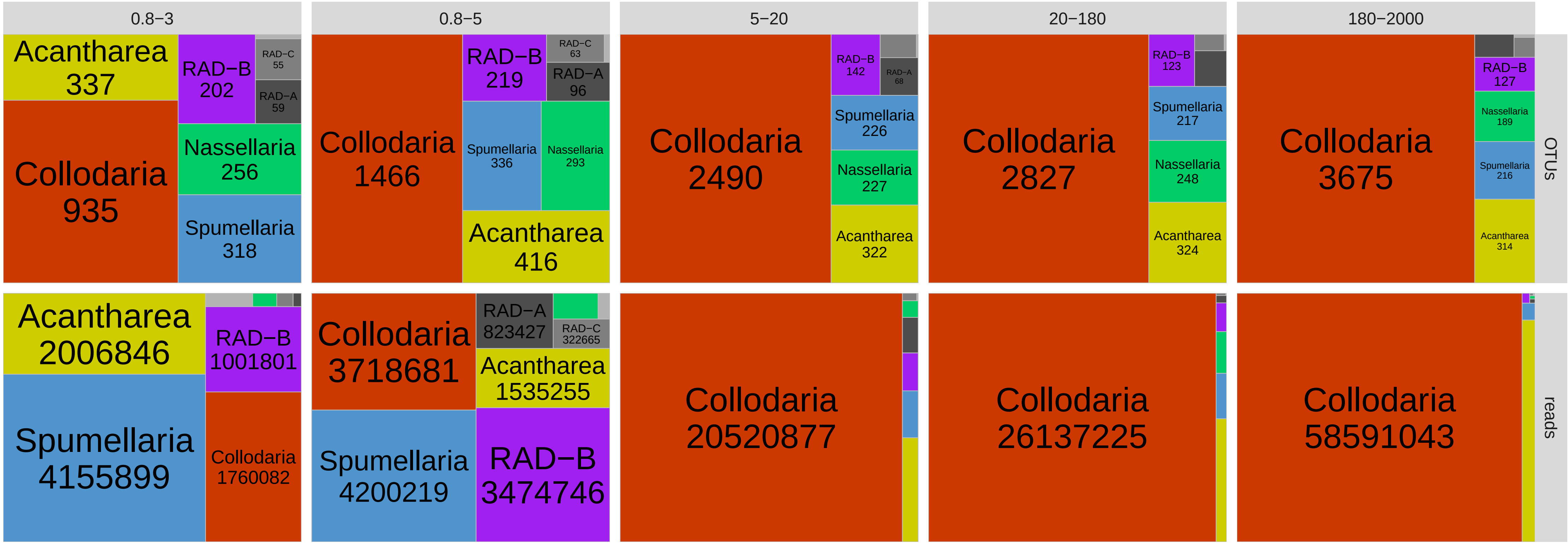

### Figure S5

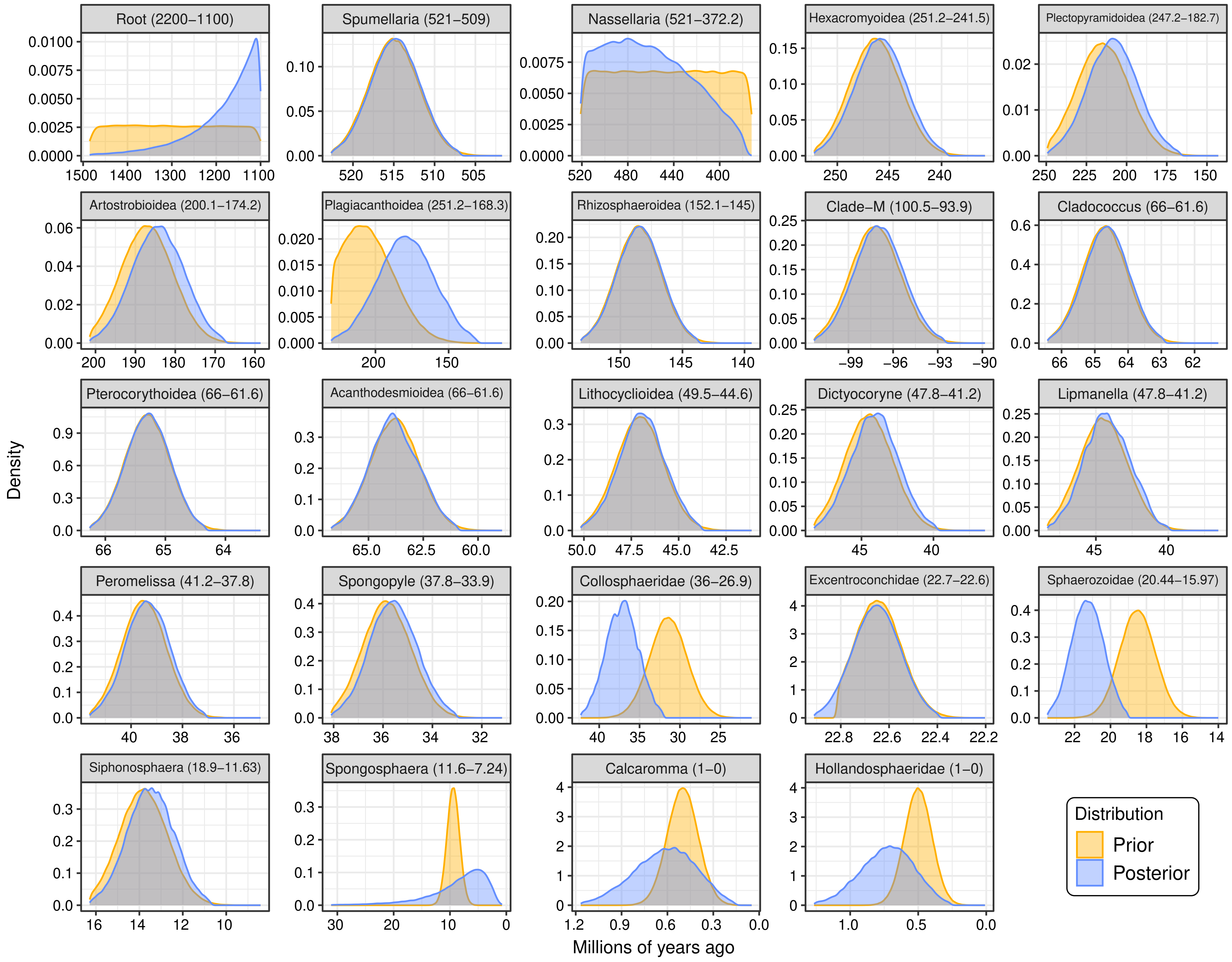

### Figure S6

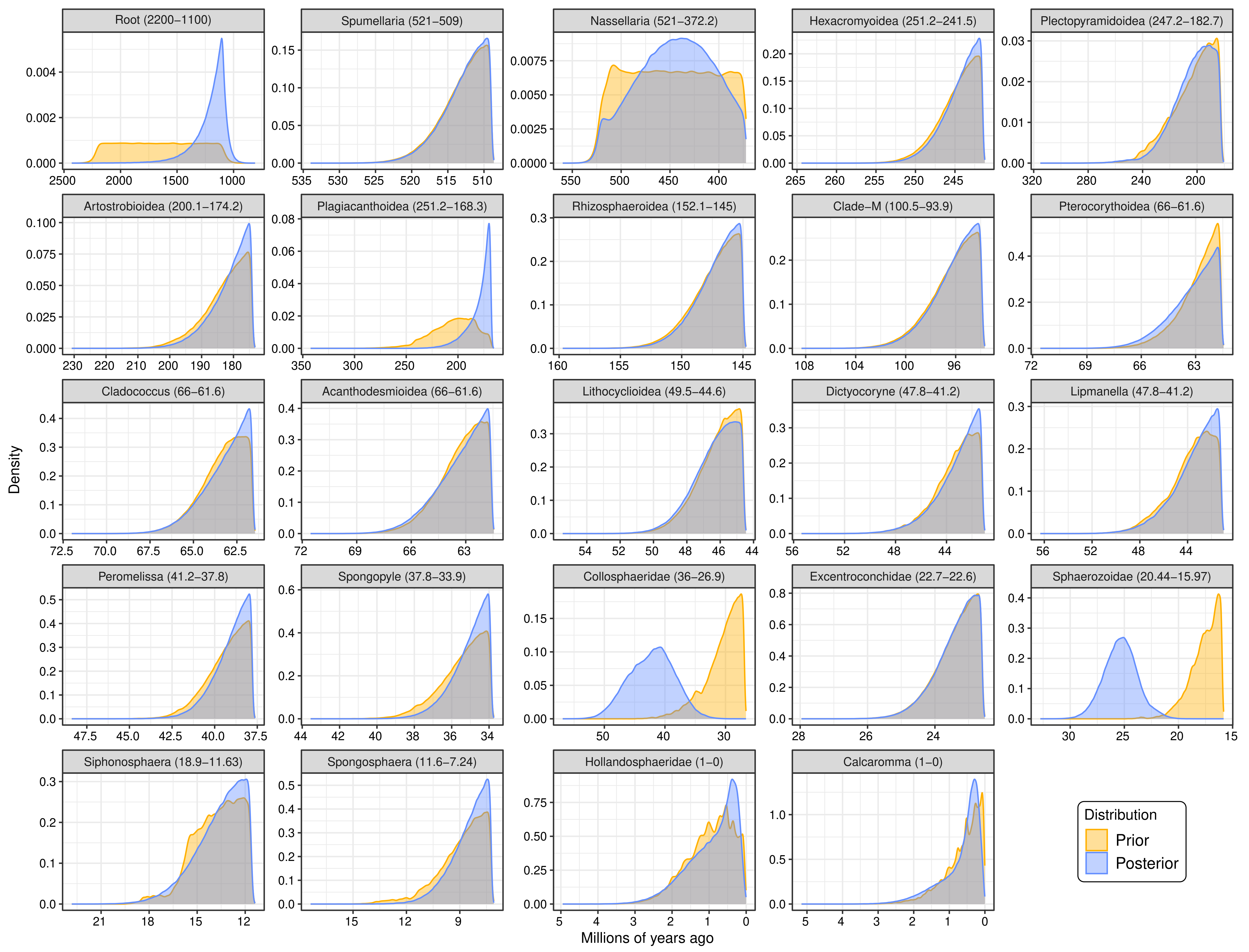

### Figure S7

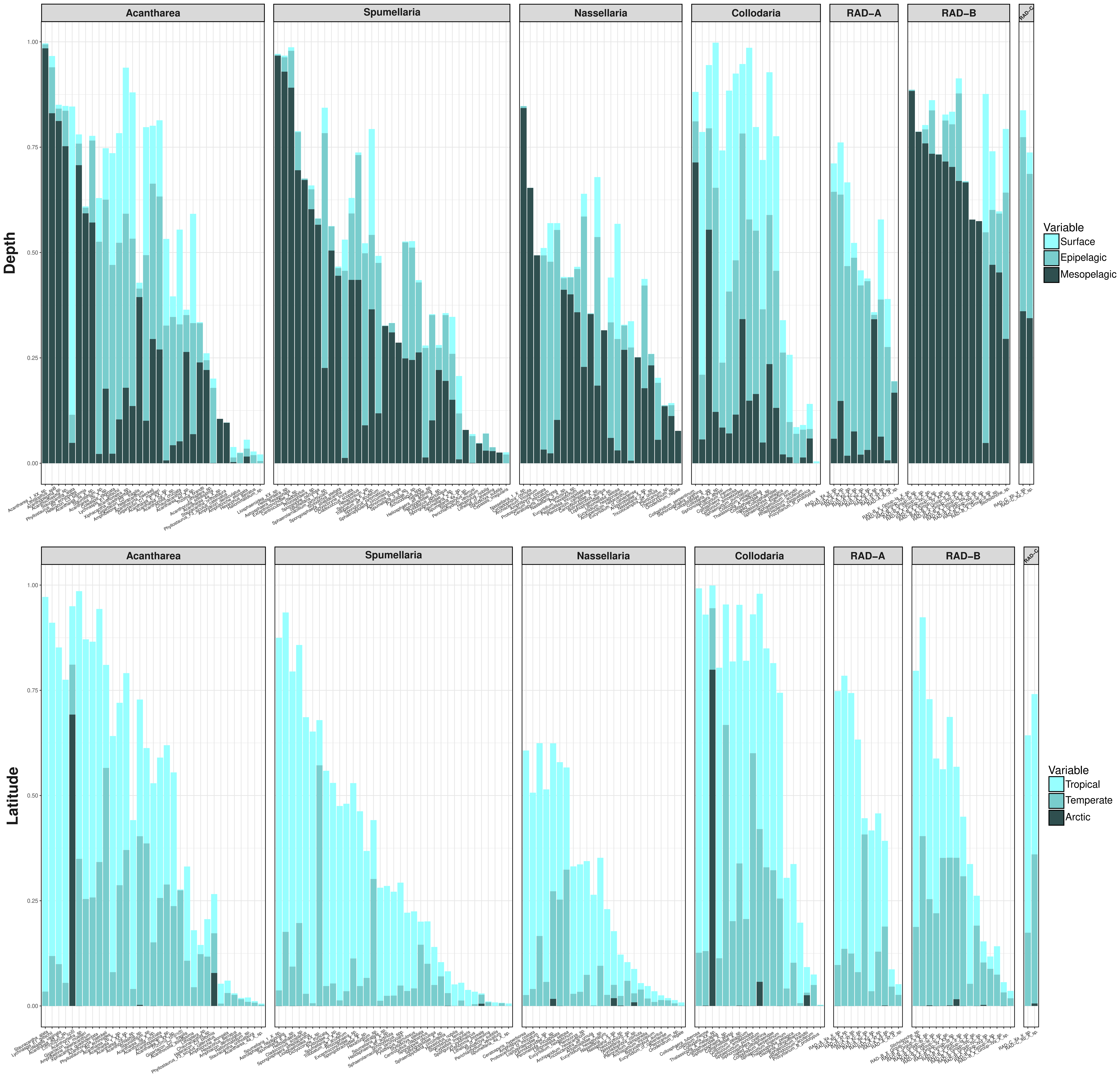
