## Supplementary material for "Diversity and evolution of Radiolaria: Beyond the stars of the ocean": File S1

### Fossil calibrations

The following lines justify the fossil calibrations chosen for the molecular clock analyses based on the minimum and maximum boundaries found in the fossil record. Unless otherwise specified, all calibrations are given first for MCMCTree followed by BEAST2 softwares. In the first software, we used skew-normal distributions in which the 97.5% probability distribution falls within the minimum and maximum boundaries (represented below as **SN[minimum-maximum]**, where **‘SN**’ is a Skew-Normal distribution). In the second software, we used normal distributions such that the 95% probability distribution falls within the minimum and maximum boundaries (represented below as **N(µ, sd)**, where ‘**N**’ is a normal distribution, ‘**µ**’ is the mean and ‘**sd**’ the standard deviation). The main reason why we have used different probability distributions in MCMCTree and BEAST2 is because they use soft and hard constraints respectively. Here we have included both Superfamily-Family and genus fossils to homogenise calibrations across the diversity and temporal axes. In addition, genus calibrations are also used due to the direct connection between extant and ancient morphologies, in an attempt to correct or adapt miscalibrations of families and/or superfamilies. Therefore, we acknowledge possible uncertainties of fossil calibrations at higher taxonomic levels (1), due to indirect connections between early fossil evidence and crown molecular clades.

**Root**: Uniform distribution [1100-2200]: The root node was calibrated between 1100 and 2200 million years ago (Ma) after the latest molecular clock results on the diversification between Alveolates and Rhizaria (2, 3) and allowing a significant uncertainty due to the lower representation of Alveolata molecular diversity in our dataset.

**Spumellaria**: SN[509-521], N(515, 3.06): Given the good agreement between the oldest radiolarian fossils with spherical forms (taxonomically classified in the extinct spumellarian genus *Paraantygopora*, early Cambrian, Series 2, ca. 521–509 Ma; (4)) and the latest molecular clock results (515 -Highest Posterior Density, HPD: 659-382- Ma; (5)).

**Nassellaria**: Uniform distribution [372.2-521]: The first appearance of Nassellaria in the fossil record is debated between primitive nassellarian forms (in the Upper Devonian) or with the first multi-segmented nassellarians (in the Early Triassic) (6, 7). Molecular clock results agree with the Devonian origin of Nassellaria (423 -HPD: 500-342- Ma; (8)), yet Collodaria was not included in the study since traditional morphology-based studies separate Collodaria as an independent order, and thus the last common ancestor of Nassellaria (as described herein) could be older.

**Hexacromyoidea group**: SN[241.5-251.2], N(246.35, 2.47): After the calibration used in (5), therein referred to as Hexastyloidea: “The family Hexastylidae [herein as Hexacaryidae] is the first representative of the superfamily Hexastyloidea [herein as Hexacromyoidea] and has its first appearance in the fossil record in the Middle Triassic (Late Anisian: ca. 246.8–241.5 Ma;(9))”. Yet, the authors pointed out a long disconnection between Triassic and Cenozoic genera belonging to this family, and recently restricted the use of the Hexacaryidae family to the Cenozoic (10). Given the paraphyletic nature of the families within Hexacromyoidea (families Hexacaryidae, Hexacromyidae and Hollandosphaeridae sharing a last common ancestor with the superfamilies Spongosphaeroidea, Spongodiscoidea, Lithocyclioidea and the environmental clade Spumellaria-1), it is possible that extant representatives diversified in the Cenozoic but have their last common ancestor in the Mesozoic. In this context, Multiarcusellidae could be a potential candidate for the last common ancestor of this group of clades with an origin in the Olenekian (247.2-251.2Ma), and would allow fossil representatives along Cretaceous and Jurassic fall within this big group filling the fossil gap.

**Plectopyramidoidea**: SN[182.7-247.2], N(214.95, 16.45): The earliest possible fossil assigned to this group is *Celluronta* Sugiyama 1997, found at the base of the Anisian (242-247.2 Ma; Middle Triassic; (11)). However, the earliest undisputed fossil assigned to Plectopyramioidea is *Cornutella* Yao 1979, found in the Pliensbachian (182.7-190.8 Ma; Lower Jurassic; (12)). In addition, the limited molecular diversity represented in our phylogenetic trees (only two specimens described as *Litharachnium* and *Polypleuris*) makes it challenging to assign the divergence of our molecular sequences to the crown Plectopyramioidea, and thus a big range is the most cautious option.

**Plagiacanthoidea**: SN[168.3-251.2], N(209.75, 21.15): The origin of this clade in the fossil record is debated between earlier representatives of the family Ximolzasinae (251.2-247.2 Ma, Olenekian, Early Triassic) and between certain representatives of the family Plagiacanthinae (168.3-170.3 Ma, Bajocian, Middle Jurassic) (9).

**Artostrobioidea**: SN[174.2-200.1], N(187.15, 6.61): After the calibration used in (8): “The genus *Artostrobium*, questionably assigned by (13), is the oldest fossil for this family, and its first appearance in the fossil record is dated in the Early Jurassic (Toarcian: ca. 183.7–174.2 Ma) (13)”. However, if we consider the genus *Ectonocorys* as an Artostrobiidae, the earliest representative was reported in the Hettangian (196.5-200.1 Ma) by (14).

**Rhizosphaeroidea**: SN[145-152.1], N(148.55, 1.81): After (5): “ The Rhizosphaeridae appeared during the late Jurassic (Tithonian: ca. 145–152.1 Ma) in the fossil record (6, 15–17)”.

**Clade-M**: SN[93.9-100.5], N(97.2, 1.68): After the calibration used in (5), therein referred to as Pylonioidea: “The family Larnacillidae are the first representatives of the superfamily Pylonioidea appearing at the beginning of the Late Cretaceous (Cenomanian: ca. 100.5–93.9 Ma; (6, 15, 18)”.

**Pterocorythoidea**: SN[61.6-66], N(65.3, 0.375): After (8): “*Cryptocarpium* is the first, but doubted, representative of this family in the fossil record and it is dated in the Late Paleocene (ca. 66–56 Ma), followed by *Podocyrtis*, the first true representative, in the transition of the Eocene (ca. 56–33 Ma) (6, 19, 20)”. In addition, *Lamptonium colymbus* (21), sample DSDP 384-11-3, 38-40 cm, is dated at 65.3 Ma according to the age model for that site in the Neptune database (22) and confidently assigned to Pterocorythoidea.

***Cladococcus*** (Cladococcoidea): SN[61.6-66], N(64.7, 0.675): The oldest occurrence corresponds to *Cladococcus nakasekoi* (21), sample DSDP 384-11-1, 38-40 cm dated at 64.7 Ma according to the age model for that site in the Neptune database (22).

**Acanthodesmioidea**: SN[61.6-66], N(63.8, 1.12): After (8): “The three families belonging to this superfamily (Acanthodesmiidae, Stephaniidae, Triospyrididae) have their first appearance in the fossil record in the Paleocene (ca. 66–56) (6, 18, 23), and the first genus described associated with these families corresponds to *Tholospyris* in the Early Paleocene (61.6-66 Ma; (24, 25))”.

**Lithocyclioidea**: SN[44.6-49.5], N(47.05, 1.25): Adapted from (5), therein referred to as Coccodiscoidea, clade E1): This family appears for the first time in the fossil record in the early middle Eocene (6, 15).

***Dictyocoryne*** (Spongodiscoidea): SN[41.2-47.8], N(44.5, 1.68): The oldest occurrence of *Dictyocoryne* is dated at base of the Lutetian (41.2-47.8 Ma, Eocene, Paleogene; (26)).

***Lipmanella*** (Plagiacanthoidea): SN[41.2-47.8], N(44.5, 1.68): The oldest occurrence of *Lipmanella* is dated at base of the Lutetian (41.2-47.8 Ma, Eocene, Paleogene; (10)).

***Peromelissa*** (Plagiacanthoidea): SN[37.8-41.2], N(39.5, 0.87): The oldest occurrence of *Peromelissa* is dated at base of the Bartonian (37.71-41.2 Ma, Eocene, Paleogene; (10)).

***Spongopyle*** (Clade-L): SN[33.9-37.8], N(35.85, 0.99)): The oldest occurrence of *Spongopyle* (27) is dated at base of the Priabonian (33.9-37.71 Ma, Eocene, Paleogene).

**Collosphaeridae**: SN[26.9-36], N(31.45, 2.32): The oldest occurrence of *Collosphaera* sp. (28), sample ODP 1218A-25X-1,30-32cm, is dated at 36 Ma according to the age model for that site in the Neptune database (22), yet the oldest occurrence of *Acrosphaera spinosa echinoides* Haeckel (in (29)), the oldest formally described species belonging to this group, Sample ODP 1218A-11H-6, 122-124 cm, is dated at 26.9 Ma according to the age model for that site in the Neptune database (22).

**Excentroconchidae**: SN[22.6-22.7], N(22.65, 0.1): First reported occurrence of *Lonchosphaera spicata* Popofsky was in sample ODP 748B-8H-6,45-47cm (30) dated at 22.6 Ma according to the age model for that site in the Neptune Database (22).

**Sphaerozoidae**: SN[15.97-20.44], N(18.5, 0.995): First reported occurrence of *Sphaerozoum punctatum* was in sample ODP 748B-7H-4,45-47cm (30) dated at 18.5 Ma according to the age model for that site in the Neptune Database (22).

***Siphonosphaera*** (Collosphaeridae): SN[11.63-18.9], N(13.82, 1.12): The oldest occurrence of *Siphonosphaera* (31) is dated at base of the Serravallian (11.63-13.82 Ma, Miocene, Neogene). However *S.*? *abelmannae* (32) has been dated at 18.9 Ma (30), although it is only questionably assigned to *Siphonosphaera*.

***Spongosphaera*** (Spongosphaeroidea): SN[7.24-11.6], N(9.42, 1.11): The oldest occurrence of *Spongosphaera* (31) is dated at base of the Tortonian (7.24-11.6 Ma, Miocene, Neogene).

***Calcaromma*** (Clade-L): SN[0-1], N(0.5, 0.25): This genus is only found in living samples (10).

**Hollandosphaeridae** (Hexastyloidea-group): SN[0-1], N(0.5, 0.25): This group is only represented by the genus *Hollandosphaera* in our phylogenetic dataset and is only found in recent sediments (33).

27.	F. Dreyer, Die Pylombildungen in vergleichend-anatomischer und entwicklungsgeschichtlicher Beziehung bei Radiolarien und bei Protisten uberhaupt, nebst System und Beschreibung neuer und der bis jetzt bekannten pylomatischen Spumellarien. *Jenaische Z. Für Naturwissenschaft Jena* **23**, 77–214 (1889).

28.	S. Funakawa, H. Nishi, Late middle Eocene to late Oligocene radiolarian biostratigraphy in the Southern Ocean (Maud Rise, ODP Leg 113, Site 689). *Mar. Micropaleontol.* **54**, 213–247 (2005).

29.	S. Kamikuri, H. Nishi, T. C. Moore, C. A. Nigrini, I. Motoyama, Radiolarian faunal turnover across the Oligocene/Miocene boundary in the equatorial Pacific Ocean. *Mar. Micropaleontol.* **57**, 74–96 (2005).

30.	S. Trubovitz, D. Lazarus, J. Renaudie, P. J. Noble, Marine plankton show threshold extinction response to Neogene climate change. *Nat. Commun.* **11**, 5069 (2020).

31.	E. Haeckel, Die Radiolarien (Rhizopoda radiaria). Eine Monographie, Bd. 1 (Text) und Bd. 2 (Atlas). (1862).

32.	J. Renaudie, D. B. Lazarus, New species of Neogene radiolarians from the Southern Ocean. *J. Micropalaeontology* **31**, 29–52 (2012).

33.	G. Deflandre, Observations et remarques sur les Radiolaires Sphaerellaires du Paléozoı̈que, à propos d’une nouvelle espèce viséenne, du genre Foremaniella Defl., parfait intermédiare entre les Périaxoplastidiés et les Pylentonémidés. *C. R. ACAD. SCI.* **VOL. 276**, 1147–1151 (1973).
